# Transcriptome-based phylogeny and whole-genome duplication in Theaceae

**DOI:** 10.1101/2021.03.26.437128

**Authors:** Qiong Zhang, Lei Zhao, Jian-Li Zhao, Ryan A. Folk, Nelson Zamora, Pamela S. Soltis, Douglas E. Soltis, Shi-Xiong Yang, Lian-Ming Gao, Hua Peng, Xiang-Qin Yu

**Affiliations:** CAS Key Laboratory for Plant Diversity and Biogeography of East Asia, Kunming Institute of Botany, Chinese Academy of Sciences, Kunming 650201, China; College of Life Sciences, University of Chinese Academy of Sciences, Beijing 100049, China; Germplasm Bank of Wild Species, Kunming Institute of Botany, Chinese Academy of Sciences, Kunming, Yunnan 650201, China; Yunnan Key Laboratory of Plant Reproductive Adaption and Evolutionary Ecology, Yunnan University, Kunming 650091, China; Department of Biological Sciences, Mississippi State University, MS 39762, United States; National Herbarium of Costa Rica (CR), Natural History Department of National Museum of Costa Rica, San José, Costa Rica; Florida Museum of Natural History, University of Florida, Gainesville, FL 32611, United States

**Author notes:** Correspondence: Xiang-Qin Yu, Hua Peng.

**Keywords:** Theaceae, phylogeny, transcriptome, low-copy nuclear genes, phylogenetic network

## Abstract

Theaceae, with three tribes and nine genera, is a family of great economic and ecological importance. Recent phylogenetic analyses based on plastid genome resolved the relationship among three tribes and the intergeneric relationships within Gordonieae and Stewartieae. However, generic level relationships within the largest tribe Theeae were not fully resolved and potential hybridization among genera within Theeae revealed previously also remains to be tested further. Here we conducted a comprehensive phylogenomic study of Theaceae based on transcriptomes and low-depth whole-genome sequencing of 57 species as well as additional plastome sequence data from previous work. Phylogenetic analyses suggested that Stewartieae was the first-diverging clade in Theaceae, consistent with previous study using plastomic data. Within Theeae, the highly supported *Apterosperma-Laplacea* clade grouped with *Pyrenaria* with maximum support based on the partitioned and unpartitioned concatenation analyses using the 610 low-copy nuclear genes, leaving *Camellia* and *Polyspora* as another sister genera in the tribe. PhyloNet analyses suggested one reticulation event within *Camellia* and *Pyrenaria* respectively, but no intergeneric reticulations were detected in Theeae. Another introgression was found between *Gordonia lasianthus* and the common ancestor of Gordonieae during the Late Oligocene. The existing land bridges (e.g. Bering land bridge) might have facilitated this ancient introgression. Further researches need to be conducted to uncover the interspecific introgression pattern within *Camellia.* Ks distribution analyses supported the tea family shared one whole-genome duplication (WGD) event Ad-β, which was recently mapped to the clade containing core Ericales, Primuloids, Polemonioids and Lecythidaceae.

## Introduction

Theaceae consists of nine genera in three tribes and contains more than 200 species of evergreen and deciduous trees and shrubs that are disjunctly distributed in temperate, subtropical and tropical areas of eastern to southeastern Asia, and eastern North America to Central and South America, i.e. an Amphi-Pacific disjunction (Kobuski, 1949; Kobuski, 1950; Prince, 1993; Stevens, 2001 onwards; Min & Bartholomew, 2007). Members of Theaceae have great economic and ecological importance, including familiar plants such as tea (e.g. *C. sinensis* (L.) Kuntze), oil plants (e.g. *C. oleifera* C. Abel) and a number of woody ornamentals (e.g. *Camellia japonica* L., *C. reticulata* Lindley), and tree representatives (e.g. *Schima)* and smaller woody lineages (e.g. *Camellia, Stewartia)* are dominant or common species of the subtropical evergreen broadleaved forests in East Asia (Tang, 2015). Due to the excessive collections and habitat destruction, several species mainly from *Camellia* have been listed as (critically) endangered, such as *C. fangchengensis* S. Ye Liang & Y. C. Zhong, *C. hekouensis* C. J. Wang & G. S. Fan, *C. piquetiana* (Pierre) Sealy (IUCN, 2020). Meanwhile, many new species (Orel, 2006; Orel & Wilson, 2010b; Orel & Wilson, 2012; Orel *et al.*, 2013; Lee & Yang, 2019; Liu *et al.*, 2019), and subgeneric taxa (Orel & Wilson, 2010a; Orel *et al.*, 2014) have been described and published for *Camellia* in the past ten years.

After the establishment of Theaceae in 1813 (Mirbel, 1813), the systematic boundaries of genera in the family as defined by morphology has changed significantly (Bentham & Hooker, 1862; Melchior, 1925; Takhtajan, 1997), from two genera to six tribes and 32 genera. Molecular systematic studies recognized three tribes and nine genera (APG I, 1998; APG II, 2003; APG III, 2009; APG IV, 2016). Since then, a number of systematic studies using DNA markers, morphology, anatomy and cytology have been conducted to explore the relationships among tribes and genera (Ye, 1990; Tsou, 1998; Prince & Parks, 2001; Yang *et al.*, 2004; Wang *et al.*, 2006; Yang *et al.*, 2006; Li *et al.*, 2013; Zhang *et al.*, 2014; Yu *et al.*, 2017b). However, many unresolved phylogenetic relationships remain in this family.

Firstly, phylogenetic relationships among the three tribes in Theaceae remain controversial. Evidence from floral development indicated a close relationship between the tribes Gordonieae and Stewartieae (Tsou, 1998; treated as subtribes in this study), which was supported by phylogenetics based on small single-copy region (SSC) of the plastome with conflicting support between non-model-based maximum-parsimony (MP), model-based maximum-likelihood (ML) and Bayesian inference (BI) analyses (Li *et al.*, 2013). However, an analysis of 46 morphological characters supported sister between Gordonieae (=Schimeae) and Theeae (Wang *et al.*, 2006), which was supported by phylogenetics of plastid *rbcL, matK* and *trnL-F,* mitochondrial *matR* and nrITS (Prince & Parks, 2001; Yang *et al.*, 2004; Yang *et al.*, 2006) and the whole plastome sequences (Yu *et al.*, 2017b). In a recently published phylogenetic context of Ericales based on 25 genomic loci from the plastid, nuclear and mitochondrial genomes, Theeae and Gordonieae were also recovered as sister with Stewartieae as the first-diverging clade (Rose *et al.*, 2018). Given conflict and uncertain support, additional nuclear genes are needed to draw a more comprehensive conclusion.

Moreover, intergeneric relationships within Theeae have been challenging to resolve, especially for the phylogenetic position of *Apterosperma* and *Laplacea.* Since the time of its original description (Chang, 1976), *Apterosperma* has been formally placed in tribe Gordonieae (=Schimeae) and considered close to *Schima* and *Franklinia* based on similar morphological characters (Ye, 1990; Chang & Ren, 1998; Tsou, 1998). A combined molecular phylogenetic analysis based on nrITS, plastid *trnL-F,* and mtDNA *matR* sequence data placed *Apterosperma* as the first-diverging lineage within Theeae (Yang *et al.*, 2004). Based on analyses of 46 morphological characters, *Apterosperma* was also placed as the first-diverging lineage within Theeae (Wang *et al.*, 2006). Using five genomic regions (chloroplast: *atpI-H, matK,psbA5’R-ALS-11F, rbcL;* nuclear: *LEAFY)* and 30 species representing four of the five genera within Theeae, Zhang *et al.* (2014) found that *Apterosperma* formed a sister relationship with *Polyspora* in the cpDNA tree, but was placed within a clade comprising *Tutcheria* (=*Pyrenaria*) and *Parapyrenaria (=Pyrenaria)* in the *LEAFY* tree. In their study, *Camellia* and *Pyrenaria* were not monophyletic, and inconsistent phylogenetic placement of some species between the nuclear and chloroplast trees were proposed to be the result of widespread hybridization among genera in Theeae. In contrast to *Apterosperma*, very few studies have included *Laplacea*. The study of Prince & Parks (2001) recovered *Laplacea* within a clade comprising *Camellia*, *Tutcheria* (=*Pyrenaria*) and *Glyptocarpa* (=*Pyrenaria*). In our previous study, *Apterosperma* and *Laplacea* were sister with strong support (MLBS=96%, BIPP=1.0). However, *Apterosperma-Laplacea* clade grouped either with *Camellia*-*Polyspora* or with *Pyrenaria* with moderate support based on different partitions of the plastome, grouped with *Polyspora* with weak support from nrITS dataset, or even as the early branching clade from the combined plastome and nrITS dateset (Yu *et al.*, 2017b).

Whole genome duplication (WGD), an important evolutionary force, was reported to occur in the common ancestor of extant seed plants, extant angiosperms and core eudicots (Ad-γ) (Jiao *et al.*, 2011; Vekemans *et al.*, 2012). WGD events were also found in the early history of numerous families such as Asteraceae, Brassicaceae, Fabaceae, Poaceae and Rosaceae (Huang *et al.*, 2016; Xiang *et al.*, 2017; Qiao *et al.*, 2019). Based on the kiwifruit *(Actinidia chinensis)* genome, researchers proposed a WGD event called Ad-β, which was shared by *Actinidia* and *Camellia* (Shi *et al.*, 2010; Huang *et al.*, 2013). However, using genome collinearity and also MAPS pipeline, Ad-β was recently mapped to the core Ericales (Leebens-Mack *et al.*, 2019), and more specifically to the clade comprising core Ericales, Primuloids, Polemonioids and Lecythidaceae (Zhang *et al.*, 2020). In Theaceae, whole genome was only available for *Camellia,* and results for WGD event from previous studies were incompatible (Xia *et al.*, 2017; Wei *et al.*, 2018; Xia *et al.*, 2020), and need more evidence and broaden sampling.

RNA-seq (transcriptomics) is becoming an important approach for plant phylogenomics, (Soltis *et al.*, 2013; Yang & Smith, 2013; Wickett *et al.*, 2014; Huang *et al.*, 2016; Yu *et al.*, 2018; Zhang *et al.*, 2020), and has been widely used to explore the origin and early diversification of land plants (Wickett *et al.*, 2014), deep-level (among eight clades) angiosperm phylogeny (Zeng *et al.*, 2014), phylogenetic relationship within eudicot, asterids and rosids (Zhao *et al.*, 2016; Zeng *et al.*, 2017; Zhang *et al.*, 2020), and intergeneric relationships for several species-rich orders or families (Caryophyllales, Asteraceae, Brassicaceae, Rosaceae and Pinaceae) (Huang *et al.*, 2015; Yang *et al.*, 2015; Huang *et al.*, 2016; Xiang *et al.*, 2017; Ran *et al.*, 2018). Additionally, the 1000 Plant Genomes Project (1KP), sequencing transcriptomes from 1124 species representing the diversity of green plants, provided resolution of much phylogenetic uncertainties across the green tree of life (Matasci *et al.*, 2014; Leebens-Mack *et al.*, 2019). Furthermore, combining evidence from plastid and nuclear genomic compartments allows the detection of cytoplasmic introgression and other forms of hybridization (Calvo *et al.*, 2013; Folk *et al.*, 2017; Guo *et al.*, 2018; Morales-Briones *et al.*, 2018; Stubbs *et al.*, 2020).

Here, transcriptomes/low-depth whole-genome sequencing of 57 species within Theaceae were sequenced, and both nuclear and plastid genes were extracted. Integrated with plastome data from our previous study, firstly we aim to reconstruct the nuclear phylogenetic framework of the tea family, to illuminate the relationships among tribes and genera and the phylogenetic position of *Apterosperma* and *Laplacea.* Furthermore, we also test the intergeneric hybridization hypothesis proposed by previous work, and investigate whether previous WGD found in Camellia was shared by other genera of Theaceae.

## Materials and Methods

### Taxon sampling, RNA sequencing and whole-genome sequencing

We collected 58 samples representing 57 species, three tribes and all nine genera of Theaceae (Table 1). Fresh, mature and healthy leaves were collected and then frozen immediately in liquid nitrogen. For some of the samples, we stored the fresh leaves in a −80°C freezer after field collection. Total RNA was extracted from the frozen leaves using the Spectrum™ Plant Total RNA Extraction Kit (Sigma). Silica-dried leaves of two species *(Stewartia malacodendron* and *Laplacea fruticosa)* were used for low-depth whole-genome sequencing (ca. 30×). RNA-Seq and low-depth whole-genome sequencing library construction, Illumina HiseqXten sequencing, raw data cleaning and quality control were performed at Novogene, China. Additionally, chloroplast genome data were obtained from our previous studies (Yu *et al.*, 2017a; Yu *et al.*, 2017b). Six species from Symplocaceae, Styracaceae, Pentaphylacaceae and Diapensiaceae were also used as outgroup taxa. The transcriptome data of these six species were downloaded from GenBank. Based on the results from previous phylogenetic studies of Theacaee, *Pyrenaria* and *Stewartia* were treated in the broad sense according to the treatment in the Flora of China (Min & Bartholomew, 2007). In our previous study (Yu *et al.*, 2017b), we found that *Laplacea grandis* grouped with *Gordonia lasianthus* and should therefore be removed into *Gordonia*, thus here we use the name *Gordonia brandegeei* H. Keng (=*Laplacea grandis*) according to Keng (1980).

**Table 1.** List of taxa sampled in this study, with voucher and Illumina reads, species names with underscore represent those species selected for PhyloNet analysis

| Taxon | Voucher specimen | Sources | No. reads (trimmed) | SRA number | Plastid genome |
| --- | --- | --- | --- | --- | --- |
| **Stewartieae** |  |  |  |  |  |
| *Stewartia calcicola* | YXQ090 | Yunnan, China | 76,314,960 | - | KY406783 |
| *Stewartia cordifolia* | YXQ144 | Guangxi, China | 101,158,084 | - | KY406775 |
| *Stewartia crassifolia* | YXQ171 | Hunan, China | 88,878,696 | - | KY406766 |
| *Stewartia malacodendron* | FLAS 260361 | Alabama, USA | 1,798,360,616 | - | KY406773 |
| *Stewartia ovata* | 18847*A | The Arnold Arboretum | 100,544,128 | - | KY406782 |
| *Stewartia pseudocamellia* | MO-6587810 | Missouri Botanical Garden | 83,044,916 | - | KY406786 |
| *Stewartia pteropetiolata* | YXQ038 | Yunnan, China | 124,092,532 | - | KY406770 |
| *Stewartia rostrata* | YXQ15072003 | Jiangxi, China | 91,741,812 | - | KY406789 |
| *Stewartia rubiginosa* | YXQ189 | Hunan, China | 91,059,484 | - | KY406777 |
| *Stewartia sinensis* | YXQ15072001 | Jiangxi, China | 91,325,600 | - | KY406748 |
| **Gordonieae** |  |  |  |  |  |
| *Franklinia alatamaha* | MO-6587811 | Missouri Botanical Garden | 101,387,448 | - | KY406774 |
| *Gordonia brandegeei* | N. Zamora 7196 | Guanacaste, Costa Rica | 104,462,244 | - | KY406761 |
| *Gordonia lasianthus* | JCRA 110687 | JC Raulston Arboretum, Raleigh, North Carolina | 82,908,172 | - | KY406790 |
| *Schima argentea* | YXQ226 | Yunnan, China | 86,065,752 | - | - |
| *Schima brevipedicellata* | YXQ072 | Yunnan, China | 89,409,700 | - | KY406784 |
| *Schima noronhai* | YXQ034 | Yunnan, China | 88,824,452 | - | KY406787 |
| *Schima sericans* | YXQ053 | Yunnan, China | 97,108,488 | - | KY406779 |
| *Schima superba* | YXQ142 | Guangxi, China | 91,704,880 | - | KY406788 |
| *Schima wallichii* | YXQ001 | Yunnan, China | 89,876,764 | - | KY406795 |
| **Theeae** |  |  |  |  |  |
| *Apterosperma oblata* | YangSX 5978 | Guangdong, China | 86,466,584 | - | - |
| *Camellia amplexifolia* | YangSX 5010 | Hainan, China | 98,672,716 | - | - |
| *Camellia assimiloides* | YangSX 5540 | Guangdong, China | 98,123,144 | - | - |
| *Camellia cordifolia* | YangSX 5551 | Guangdong, China | 96,616,940 | - | - |
| *Camellia cuspidata* | YangSX 5118 | Hubei, China | 78,369,336 | - | - |
| *Camellia flavida* | YangSX 5865 | Guangxi, China | 92,242,904 | - | - |
| *Camellia fluviatilis* | YangSX 4033 | Guangxi, China | 68,780,488 | - | - |
| *Camellia grijsii* | YangSX 6064 | Guizhou, China | 120,959,328 | - | - |
| *Camellia gymnogyna* | YangSX 5953 | Guangxi, China | 94,768,804 | - | - |
| *Camellia huana* | YangSX 5653 | Guizhou, China | 111,316,580 | - | - |
| *Camellia ilicifolia* | YangSX 5287 | Guizhou, China | 95,559,812 | - | - |
| *Camellia longipedicellata* | YangSX 5926 | Guangxi, China | 90,347,792 | - | - |
| *Camellia longissima* | YangSX 5079 | Guangxi, China | 99,050,172 | - | - |
| *Camellia luteoflora* | YangSX 6063 | Guizhou, China | 123,788,308 | - | - |
| *Camellia pilosperma* | YangSX 4714 | Guangxi, China | 92,105,712 | - | - |
| *Camellia pitardii* var. *compressa* | YangSX 4576 | Hunan, China | 76,531,640 | - | - |
| *Camellia semiserrata* | YangSX 5555 | Guangdong, China | 86,751,660 | - | - |
| *Camellia sinensis* var. *pubilimba* | YangSX 5927 | Guangxi, China | 85,785,132 | - | - |
| *Camellia szechuanensis* | YangSX 5064 | Sichuan, China | 88,726,296 | - | - |
| *Camellia tsingpiensis* | YangSX 5798 | Guangxi, China | 90,514,420 | - | - |
| *Camellia tuberculata* | YangSX 5202 | Chongqing, China | 90,076,900 | - | - |
| *Laplacea fruticosa* | NZ10477 | Puntarenas, Costa Rica | 1,532,313,620 | - | - |
| *Polyspora axillaris* | YXQ099 | Hainan, China | 92,917,500 | - | KY406760 |
| *Polyspora chrysandra* | YXQ221 | Yunnan, China | 84,011,964 | - | - |
| *Polyspora dalgleishiana* | BROWP 501 | Royal Botanic Garden<br>Edinburgh | 83,838,300 | - | KY406769 |
| *Polyspora hainanensis* | YXQ097 | Hainan, China | 97,170,208 | - | KY406776 |
| *Polyspora longicarpa* | YangSX 4779 | Yunnan, China | 89,251,900 | - | KY406768 |
| *Polyspora speciosa* | YXQ145 | Guangxi, China | 96,508,612 | - | KY406754 |
| *Pyrenaria hirta* var. *cordatula* | YXQ169 | Guangxi, China | 92,426,012 | - | KY406785 |
| *Pyrenaria hirta* var. *hirta* | YangSX 4067 | Guangxi, China | 100,342,800 | - | - |
| *Pyrenaria jonquieriana*<br>subsp. *multisepala* | YXQ106 | Hainan, China | 87,583,392 | - | KY406772 |
| *Pyrenaria khasiana* | YangSX 5046 | Myanmar, Kachin | 88,528,812 | - | KY406756 |
| *Pyrenaria menglaensis* | YXQ211 | Yunnan, China | 87,085,244 | - | KY406747 |
| *Pyrenaria microcarpa* var. *microcarpa* | YXQ101 | Hainan, China | 81,080,788 | - | KY406764 |
| *Pyrenaria oblongicarpa* | YXQ216 | Yunnan, China | 98,658,408 | - | KY406781 |
| *Pyrenaria pingpienensis* | YXQ210 | Yunnan, China | 96,601,500 | - | - |
| *Pyrenaria shinkoensis* | YangSX 5038 | Taiwan, China | 101,120,720 | - | - |
| *Pyrenaria spectabilis* var. *greeniae* | YXQ172 | Hunan, China | 93,451,072 | - | KY406753 |
| *Pyrenaria spectabilis* var. *spectabilis* | YXQ155 | Guangxi, China | 88,012,020 | - | KY406765 |
| **Outgroup** |  |  |  |  |  |
| *Eurya acuminatissima* | - | - | 25,009,135 | SRX2786652 | - |
| *Galax urceolata* | - | - | 9,647,946 | ERX2099546 | - |
| *Sinojackia xylocarpa* | - | - | 10,611,525 | ERX2099565 | - |
| *Symplocos tinctoria* | - | - | 10,196,542 | ERX2099566 | - |
| *Symplocos paniculata* | - | - | 28,374,406 | SRX1601992 | - |
| *Ternstroemia gymnanthera* | - | - | 14,421,549 | ERX2099558 | - |

-represents the chloroplast genes were extracted from the transcriptome data or no data were available.

### Sequence assembly and ortholog identification

Quality control of the raw sequencing reads was performed using the Fastp version 0.20.1, adapter, reads containing N and reads with low quality score (percentage of base <=Q20) were removed. Trinity v2.8.4 was used to conduct *de novo* assembly of cleaned Illumina RNA-Seq reads of each species (Grabherr *et al.*, 2011; Haas *et al.*, 2013). TransDecoder v5.3.0 was used to identify candidate coding regions with default parameters. ORFs (open reading frames) with a minimum length of 100 amino acids (AA) were used for further analyses. CD-HIT v4.6 (Li & Godzik, 2006) was used to remove redundant contigs with a threshold of 0.99. Orthogroups (OG) were identified and filtered following the pipelines proposed by Yang & Smith (2014). For the data obtained from low-depth whole-genome sequencing, we first *de novo* assembled the genome using Platanus (http://platanus.bio.titech.ac.jp/platanus-assembler/platanus-1-2-4) with default parameters. Then RepeatMasker and RepeatModeler (http://www.repeatmasker.org/) were performed to identify tandem repeats and TEs. We carried out gene annotation using de novo gene prediction (AUGUSTUS, http://evidencemodeler.github.io/) and homolog prediction (Exonerate, https://www.ebi.ac.uk/about/vertebrate-genomics/software/exonerate), then generated an integrated gene set using EVidenceModeler (http://evidencemodeler.github.io/). Preliminary gene trees were reconstructed using RAxML v8.2.12 (Stamatakis, 2014). To reduce potentially misidentified orthologs, we examined single-gene trees of the 631 OGs and found three tribes within Theaceae was not monophyletic in 21 gene trees. Because all three tribes within Theaceae was consistently monophyletic in previous studies (Prince & Parks, 2001; Yang *et al.*, 2004; Li *et al.*, 2013; Yu *et al.*, 2017b), these 21 OGs likely contain hidden paralogs and are therefore problematic for downstream analysis. Therefore, we excluded the 21 OGs to yield a final gene data set with 610 OGs (hereafter refer to the reduced 610 OGs dataset). For those species without chloroplast genome data, we first assembled the chloroplast genome using GetOrganelle v1.6.2e (Jin *et al.*, 2020) and the protein-coding sequences (CDS) were extracted from the transcriptome and whole-genome sequencing data. For those species with already complete chloroplast genomes, protein-coding sequences were extracted following parallel methods to yield a combined matrix.

### Phylogenetic analyses and evolutionary network

The obtained nucleotide sequences were aligned with MAFFT v7.407 (Katoh & Standley, 2013). Alignment statistics were calculated by AMAS (Borowiec, 2016). Both concatenation and coalescent methods were used to reconstruct intergeneric relationships of Theaceae. For concatenation, partitioned and unpartitioned Maximum-Likelihood analyses and Bayesian inference (BI) were performed by using RAxML v8.2.12 (Stamatakis, 2014) and MrBayes v3.2.6 (Ronquist *et al.*, 2012). In the unpartitioned ML analysis, GTRGAMMA model was used and bootstrap support (BS) value were calculated using 1000 replicates. In the unpartitioned BI analysis, parameters were set to nst=6 and rates=gamma. Four chains were run for 2,000,000 generations with random initial trees, every 100 generations were sampled and the first 25% of the samples were discarded as burn-in. In the partitioned analyses, partitioning schemes and models were selected by using PartitionFinder version 2.1.1 for 610 OGs (Lanfear *et al.*, 2016) and 188 subsets were obtained for the CDS dataset. For coalescent analysis, a ML gene tree was reconstructed for each LCO with RAxML using the same parameter settings as above. The best ML gene trees and 100 bootstrap replicate trees generated from each LCO were used to estimate species tree and supporting values in ASTRAL v5.6.3 (Mirarab *et al.*, 2014). Support of the ASTRAL-II species tree was quantified using the local posterior probability (LPP) of a branch as a function of its normalized quartet support (Sayyari & Mirarab, 2016). We used PhyParts (Smith *et al.*, 2015) to examine patterns of gene tree concordance and conflict within the nuclear genome and to reveal subsets of the nuclear genome supporting alternative relationships in the plastid topology, visualizing the results using the program phypartspi echarts (https://github.com/mossmatters/phyloscripts/). To test the previous hypothesis of intergeneric hybridization, we used PhyloNet v3.8.0 (Than *et al.*, 2008) to infer an evolutionary network for Theaceae, using the command “InferNetwork_MPL” under a maximum pseudo-likelihood framework (Yu & Nakhleh, 2015) and 505 individual gene trees. Given that the computational time needed by network methods scales very rapidly with taxon number (Folk *et al.*, 2018), we reduced the sampling to a computationally tractable size (22 species in total, fewer than 30 taxa), taking one (i.e. *Apterosperma*, *Franklinia* and *Laplacea*) to six (i.e. *Camellia*) species of each genus, and only one outgroup species (different outgroup species were selected for each gene tree because of missing data) (Table 2). The maximum pseudo-likelihood algorithm requires *a priori* specification of the number of reticulating branches; the number of reticulations were set as one, two, three and four in repeated analyses per developer recommendations. Gene trees with branches <50% BS were collapsed, and five optimal networks returned for each analysis. The command CalGTProb was used to compute the likelihood scores and select the best network.

**Table 2.** Total log probability of each phylogenetic network

| Reticulations | Total log probability |
| --- | --- |
| 1 | -380.3215 |
| 2 | -380.3215 |
| 3 | -380.1597 |
| 4 | -379.6899 |

### Whole genome duplication analysis

In order to investigate the ancient whole-genome duplications (WGD) in Theaceae, we applied the Python package “wgd” (Zwaenepoel & Van de Peer, 2019) to construct the synonymous substitutions (*K*s) distributions (ranging from 0.05 to 3) among paralogs from 56 Theaceae transcriptomes and six outgroup transcriptomes. Using the command “mcl” to blast and cluster sequences with each CDS, the command “ksd” and “mix” was used to construct the Ks distribution and mixture modeling of *K*s distributions, respectively. For the analysis of the mixture model, we chose the method BGMM in wgd package.

## Results

### Characteristics of transcriptomes and datasets

We sequenced 5.85 to 10.6 Gb of transcriptome data from 56 individuals of 55 species, and 138.7 and 165.7 Gb whole-genome sequencing data for each of *Laplacea fruticosa* and *Stewartia malacodendron.* In total, our phylogenetic analysis represented 58 individuals of 57 species from all nine genera and three tribes of Theaceae (Table 1). To construct the plastid matrix, plastid protein-coding genes from 27 species were extracted from the transcriptome or whole-genome sequencing data generated here, while plastid genome from 31 species were obtained from complete chloroplast genomes already available from our previous study. In total, 610 OGs were obtained from 56 transcriptomes and two whole-genome sequencing datasets, with the aligned length ranging from 309 bp to 8,854 bp and missing data percentage from 0% to 29.65%. The aligned length of the concatenated 610 OGs was 858,606 bp, with 227,708 (26.5%) variable sites, 117,025 (13.6%) parsimony informative sites and 21.67% missing data. The alignment length of the concatenated 80 plastid coding genes was 69,225 bp, with 7,236 (10.5%) variable sites, 3,695 (5.3%) parsimony informative sites and 11.88% missing data.

### Phylogenetic relationships and network within Theaceae

The topology recovered from both partitioned and unpartitioned RAxML analyses based on the concatenated 610 low-copy nuclear genes strongly supported the sister relationship between Theeae and Gordonieae (MLBS=100%, PP=1.00, Fig 1, Fig S1). Additionally, based on Phyparts analysis of 509 low-copy nuclear genes, 408 gene trees (80.2%) supported the above topology (Fig. 2). Coalescent-based species tree inferred from ASTRAL yielded a concordant relationship among the three tribes with the concatenation analysis (LPP=1.00, Fig S3). RAxML analyses based on the 80 protein-coding genes of the plastid genome also supported a sister relationship between Theeae and Gordonieae (MLBS=100%, PP=1.00, Fig S3). Hence support was strong and unanimous across data partitions and analytical methods employed here.

**Figure 1.**
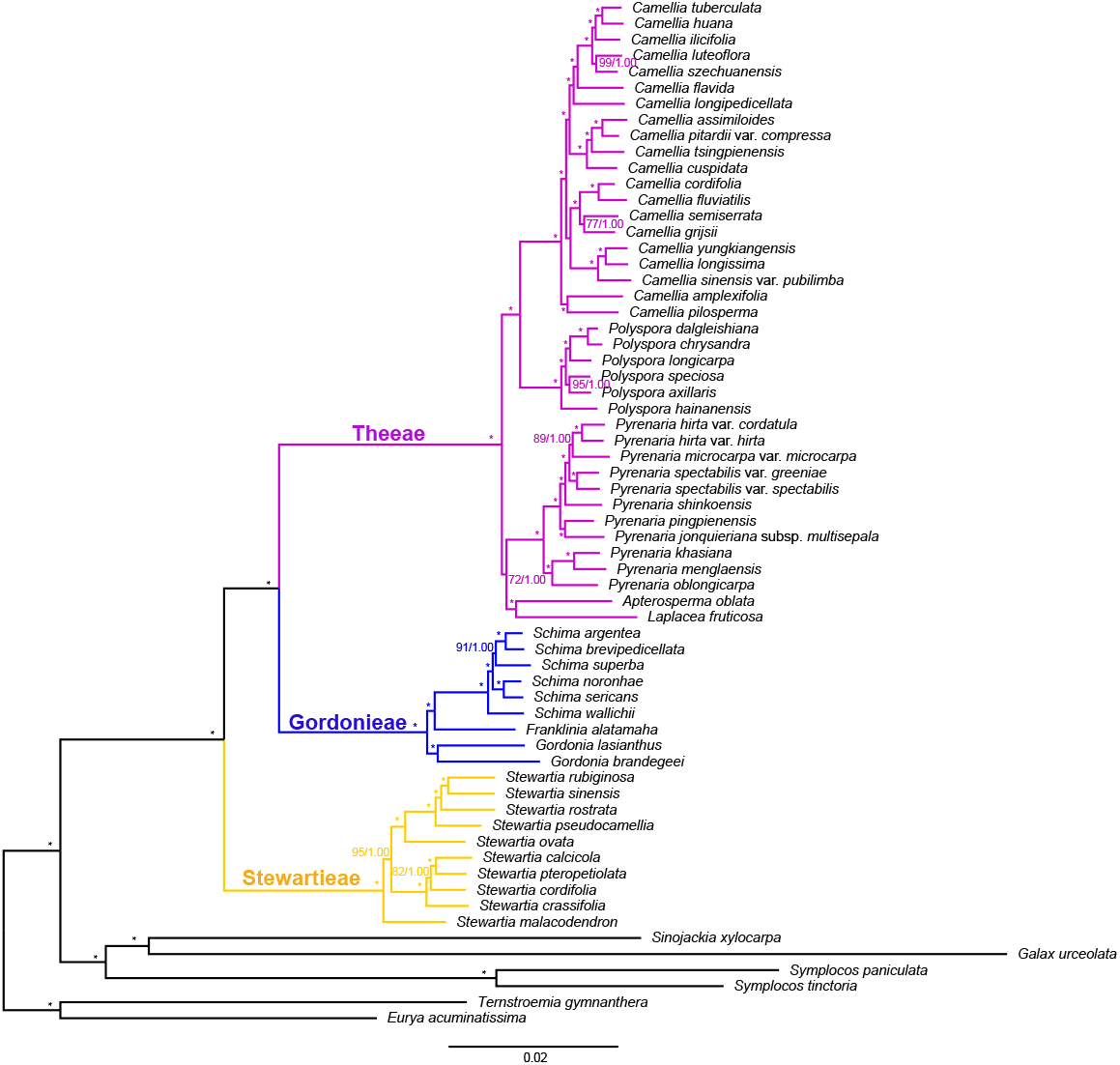
Phylogeny of Theaceae inferred from partitioned maximum likelihood (ML) analysis of the concatenated 610 low-copy nuclear genes, numbers associated with nodes indicate ML bootstrap support (BS)/Bayesian inference (BI) posterior probability (PP) values. Asterisks represent nodes with 100% support from both analyses.

**Figure 2.**
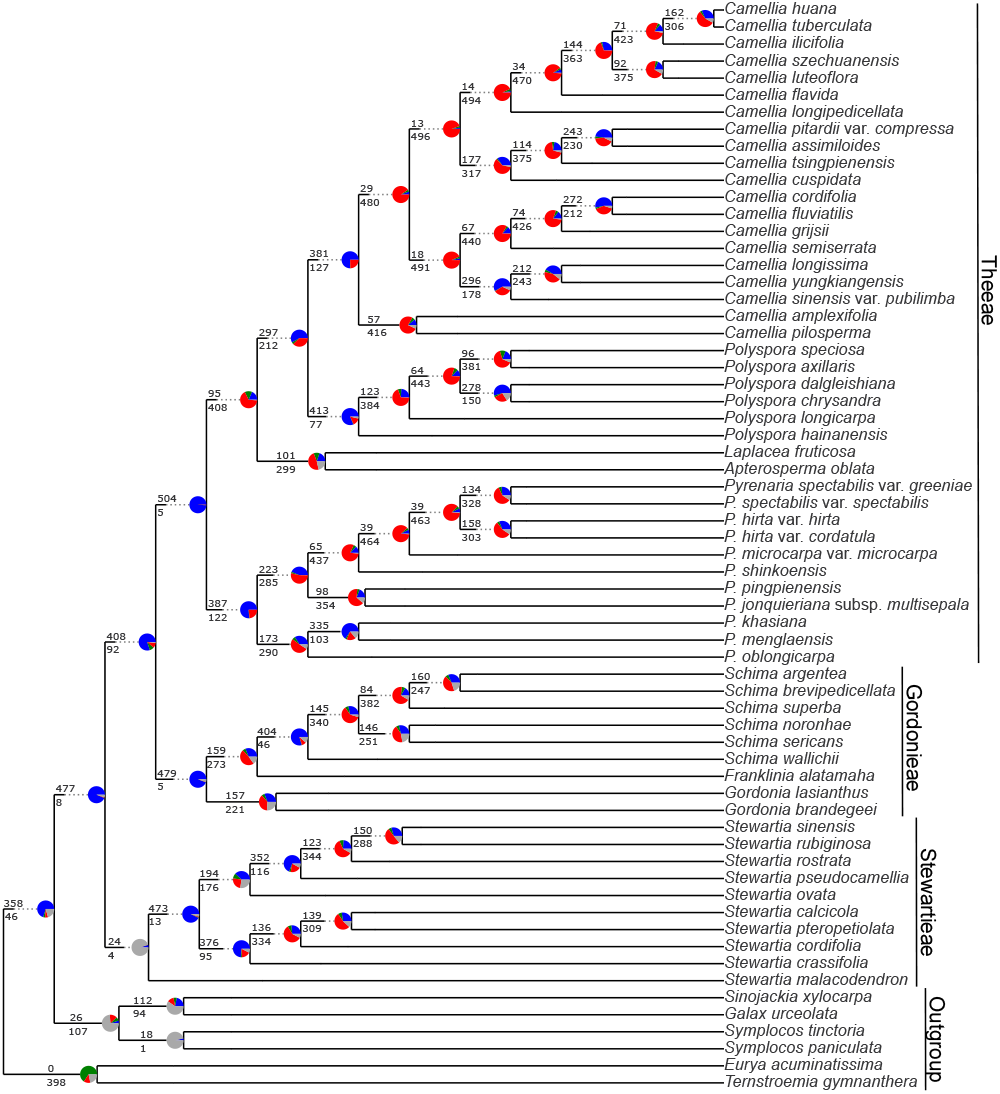
Patterns of gene-tree concordance and conflict of Theaceae based on the PhyParts analysis. The tree topology used was that inferred by ASTRAL. The pie charts at each node show the proportion of genes in concordance (blue), conflict (green = a single dominant alternative; red = all other conflicting trees), and without enough information (gray). The numbers above and below each branch are the numbers of concordant and conflicting genes at each bipartition, respectively.

Generic level relationships recovered within Gordonieae and Stewartieae were largely consistent with previous phylogenetic studies of Theaceae. Within Theeae, the sister relationships between *Camellia* and *Polyspora* and between *Apterosperma* and *Laplacea fruticosa* were both maximumly supported in all of the analyses using trascriptomic data (Fig. 1, Fig. 2, Fig. S2, Fig. S3). Nevertheless, the position of the *Apterosperma-Laplacea* clade changed between different analyses. In both of the partitioned and unpartitioned concatenation analyses using the 610 low-copy nuclear genes, the *Apterosperma-Laplacea* clade grouped with *Pyrenaria* with moderate support (MLBS=72%, PP=1.00; Fig. 1; MLBS=67%, PP=1.00; Fig. S2). The ASTRAL topology (Fig. S3), while largely congruent overall with the results from concatenated analyses (Fig. 1, Fig. S2), indicated that *Apterosperma-Laplacea* clade was weakly supported as sister to *Camellia-Polyspora* clade (LPP=0.38, Fig. S3). At lower taxonomic levels, PhyParts recovered strong discordance across many parts of the tree, with only 95 gene trees supporting the sister relationship between the *Apterosperma-Laplacea* clade and the *Camellia-Polyspora* clade, and 408 gene trees support conflicting/alternative resolutions (Fig. 2). Phylogenetic trees based on the plastid 80 protein-coding genes dataset were highly consistent with our previous study with the exception that *Apterosperma* was sister to *Camellia* (MLBS=86%, PP=1.00, Fig. S1), and *Laplacea fruticosa* grouped with *Pyrenaria* with weak support (MLBS=62%, PP=0.93, Fig. S1).

For the PhyloNet analyses, the inferred network with the highest log pseudo-likelihood (−379.6899) included three reticulations (Table 2; Fig. 3). This analysis suggested one reticulation for the clade comprising *Pyrenaria spectabilis* var. *spectabilis and Pyrenaria jonquieriana* subsp. *multisepala,* with contribution from *Pyrenaria oblongicarpa* and the ancestor of *Pyrenaria. Camellia tsingpienensis* was recovered as a hybrid between *Camellia fluviatilis and Camellia huana.* It also recovered a reticulation event suggesting that *Gordonia brandegeei* (previously difficult to place and treated as a species of *Laplacea)* descends from *Gordonia lasianthus* and the common ancestor of *Gordonia, Franklinia,* and *Schima* (i.e. Gordonieae). All other analyses (reticulation numbers =1, 2, 3), for which likelihood was suboptimal, only detected intergeneric reticulation within *Camellia* (Fig. 3).

**Figure 3.**
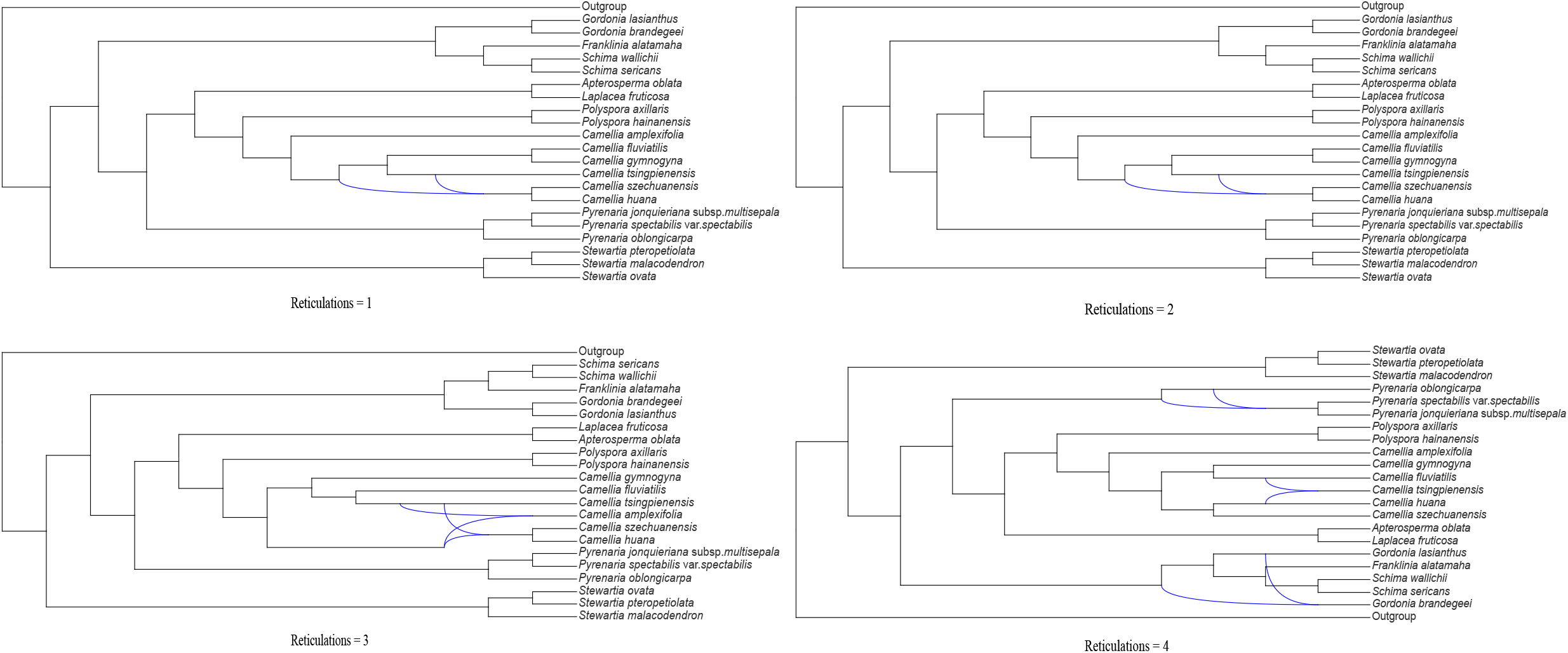
The optimal phylogenetic network of Theaceae inferred using PhyloNet, with the number of reticulations as one, two, three and four. The scenario with four reticulations had the best log pseudo-likelihood.

### Whole genome duplication

*K*_S_ (synonymous substitution rate) analyses using all 56 ingroup transcriptomes suggested that all except one species from Theaceae presented a peak at around 0.4 (Fig. 4, Fig. S4), which is consistent with one of the WGD (*Ks*=0.36) reported in tea tree *(Camellia sinensis* var. *sinensis)* genome. No *Ks* peak was found in *Stewartia ovata*, this might be due to the stochastic error, or because the transcriptome data only represent the expressed mRNA in the tissues (e.g. roots, leaves, flowers) sampled. For six outgroup species, consistent *Ks* peaks were found at around 0.4 (Figure S4), indicating Symplocaceae, Styracaceae, Pentaphylacaceae and Diapensiaceae shared this WGD event (i.e. Ad-β) with Theaceae.

**Figure 4.**
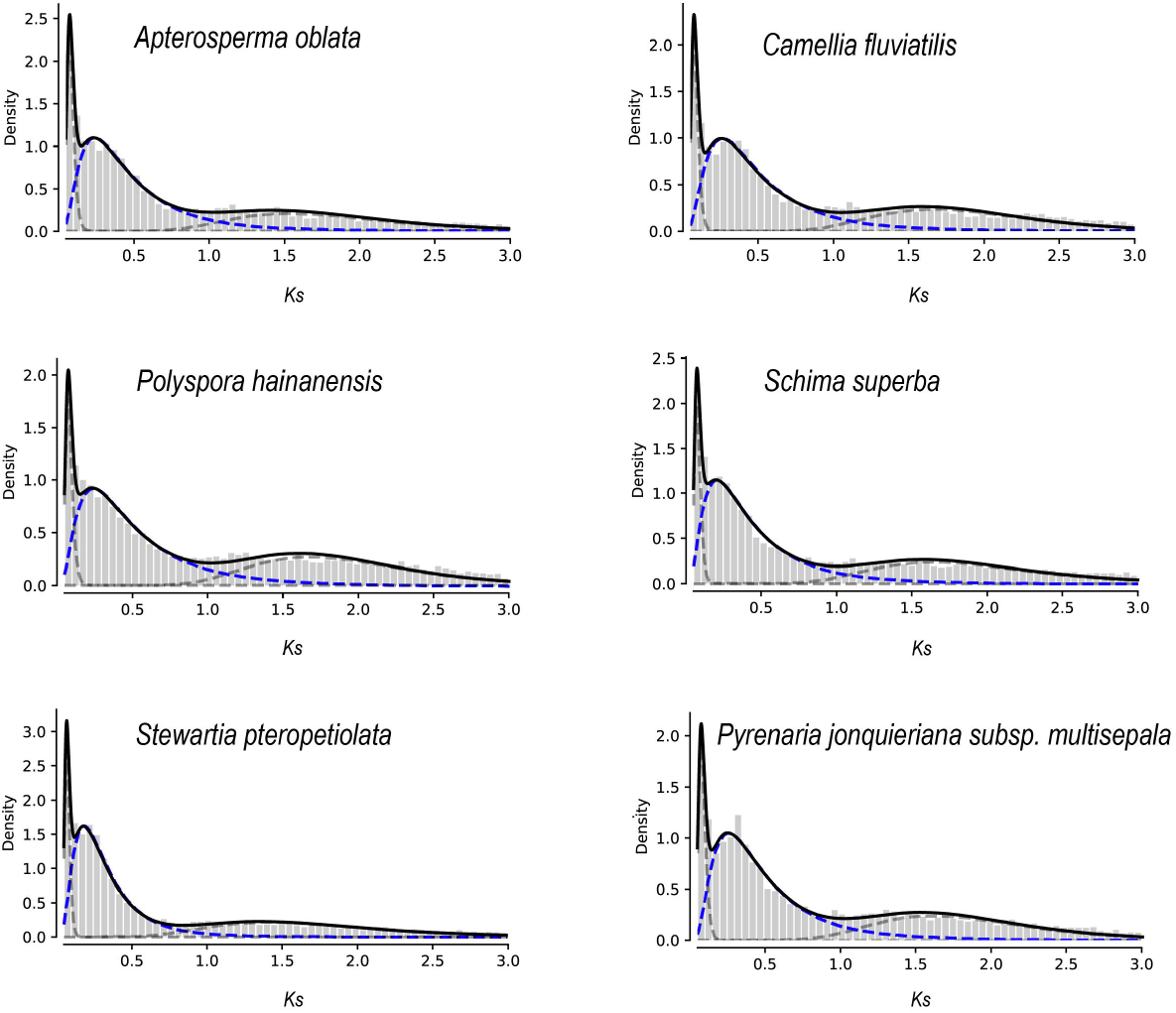
*Ks* distribution for paralogs with mixture models of inferred WGDs (blue dashed line) for *Apterosperma oblata*, *Camellia fluviatilis*, *Polyspora hainanensis*, *Schima superba*, *Stewartia pteropetiolata and Pyrenaria jonquieriana* subsp. *multisepala.*

## Discussion

### Relationships among tribes

Previous phylogenetic studies have not been able to conclusively elucidate the relationships among the three tribes within Theaceae, due in part to incomplete taxon sampling and few loci, whether based on a few plastid and nuclear loci or half of the SSC of the plastid genome (Prince & Parks, 2001; Yang *et al.*, 2004; Yang *et al.*, 2006; Li *et al.*, 2013). Using 25 genomic loci from plastid, nuclear and mitochondrial genome from 4,531 species from Ericales, Theeae and Gordonieae grouped together (MLBS>70%, PP>0.95) with Stewartieae as the first-diverging clade in Theaceae, but only 10 species were included in their study (Rose *et al.*, 2018). Our previous study supported a sister relationship between Theeae and Gordonieae (MLBS=91%, PP=1.00) based on the combined plastome and nuclear ribosomal DNA dataset (Yu *et al.*, 2017b). Here we present the phylogenetic framework of Theaceae using 610 orthologous low-copy nuclear genes. The obtained topology from both of the concatenation and coalescence analyses of the 610 low-copy nuclear genes consistently supported a sister relationship between Theeae and Gordonieae (MLBS=100%, PP=1.00, LPP=1.00, Fig. 1, Fig. S2), consistent with the results from plastid genome (MLBS=91%, PP=1.00, Fig. S3). In addition, 408 among 509 low-copy nuclear gene trees from Phyparts analysis supported the above topology (Fig. 2). The ((Gordonieae, Theeae) Stewartieae) relationship was consistent with the evolutionary pattern of the endosperm in Theaceae, as discussed in our previous study (Yu *et al.*, 2017b).

### Intergeneric relationships

The intergeneric relationships within Gordonieae and Stewartieae have been fully resolved in previous studies (Prince & Parks, 2001; Yang *et al.*, 2004; Yang *et al.*, 2006; Li *et al.*, 2013; Yu *et al.*, 2017b). However, the relationships among the five genera in Theeae have been controversial. *Laplacea* falled into the clade comprising *Camellia, Tutcheria (=Pyrenaria)* and *Glyptocarpa* (=*Pyrenaria*) (Prince & Parks, 2001). Zhang *et al.* (2014) revealed that *Apterosperma* formed a sister relationship with *Polyspora* (MLBS=73%, PP=1.00) in the cpDNA tree, but these two genera were placed in a clade comprising *Tutcheria* (=*Pyrenaria*) and *Parapyrenaria (=Pyrenaria)* (MLBS=68%, PP=0.72) in the *LEAFY* tree. Based on the 610 low-copy nuclear genes, the resolution of the relationships among the five genera in Theeae has been substantially improved. the *Apterosperma*-*Laplacea* clade received maximum support in both of the partitioned and unpartitioned concatenation analyses using the 610 low-copy nuclear genes, and grouped with *Pyrenaria* with 100% support (Fig. 1, Fig. S1). Although the ASTRAL topology suggested *Apterosperma-Laplacea* clade was weakly supported as sister to *Camellia-Polyspora* clade (LPP=0.38, Fig. S2), only 95 out of 509 nuclear genes supported this topology (Fig. 2). The strongly supported *Apterosperma*-*Laplacea* clade also grouped with *Pyrenaria* with moderate support based on the whole plastid genome dataset (MLBS=67%), the SSC (small single-copy region, MLBS=80%) dataset and the protein-coding gene dataset (MLBS=75%) from our previous study (Yu *et al.*, 2017b). Taken the evidence from plastid genome and trascriptome data, we suggest (((*Apterosperma-Laplacea*)*, Pyrenaria*), (*Camellia-Polyspora*)) as the most likely topology.

### Phylogenetic network inference

In the study of Zhang *et al.* (2014), *Camellia* and *Pyrenaria* were not monophyletic, and widespread hybridization among genera in Theeae was proposed. Phylogenetic conflicts found in *Stewartia* were also suggested to be caused by ancient introgressive hybridization following species diversification, leading to diverging histories in the nuclear and plastid genomes (Lin *et al.*, 2019). However, while our PhyloNet analyses supported the presence of hybridization in the history of Theaceae, it did not support the specific reticulation scenario suggested by Zhang *et al.* (2014); *Camellia* and *Pyrenaria* were supported as monophyletic based on the 610 low-copy nuclear genes (Fig. 1, Fig. S1). The best fit network, with three reticulation events in total, suggested two reticulation events within *Camellia* and *Pyrenaria* respectively (but no intergeneric reticulations in Theeae), and another between *Gordonia lasianthus* and the common ancestor of *Gordonia*, *Franklinia*, and *Schima* (i.e. Gordonieae). One possibility for disagreement between our work and studies could be species misidentification in the study of Zhang *et al.* (2014), but further work is needed to address hybridization more comprehensively.

All other PhyloNet analyses (reticulations=1, 2, 3) had aspects similar to the full four-reticulation analyses, suggests a clear pattern of intrageneric gene flow within *Camellia.*

Under the favored scenario involving Gordonieae, this would suggest a minimum date of reticulation during Late Oligocene, ca. 26.2 million years ago (Ma; 95%HPD=23.3–32.2 Ma) given a recent molecular dating analysis of Theaceae (Yu *et al.*, 2017b). This is a plausible scenario as land bridges (e.g. Bering land bridge) existed during Late Oligocene, the eastern Asia and eastern North American flora was likely continuous across high latitude of Northern hemisphere (Tiffney, 1985; Tiffney & Manchester, 2001; Milne, 2006), allowing for species contact and opportunities for hybridization. Fossils of Theaceae are known from high latitude Northern hemisphere localities such as North America and Germany during Ecocene and Oligocene (Grote & Dilcher, 1992; Kvacek & Walther, 1998; Wilde & Frankenhauser, 1998; Kvaček, 2004). Overall, our work, while differing from previous studies in the specific scenario, supports ancient introgression at tribe level within Theaceae that is consistent with biogeographic patterns of the group.

Previous studies have likewise found evidence of hybridization event in the genus. Firstly, under artificial conditions, cultivated ornamental camellias descending from hybridization have been widely used in horticulture (Nishimoto *et al.*, 2003; Tanaka *et al.*, 2005; Xu *et al.*, 2018). Secondly, under natural conditions, Cambod tea (cultivated tea of *C. sinensis* var. *assamica)* was suggested to have originated through hybridization between different tea types (Meegahakumbura *et al.*, 2016). Gene introgression was also detected between the cultivated *C. sinensis* var. *assamica* and *C. taliensis*, and *C.taliensis* has been suggested to be genetically involved in the domestication of *C.sinensis* var. *assamica* (Li *et al.*, 2015). All of our PhyloNet analyses consistently supported reticulation events within *Camellia* (Fig. 3). For the best-fit scenario, *Camellia tsingpienensis* (Guangxi and SE Yunnan in China, Northern Vietnam) was recovered as a hybrid between *Camellia fluviatilis* (Guangdong, Guangxi and Hainan in China, Northeastern India, Northern Myanmar) and *Camellia huana* (Guangxi and Guizhou in China). The two putative parents show distribution overlap in Guangxi province of China, thus interspecies gene flow is likely to occur under natural conditions. Further work with increased taxon sampling will need to be conducted to uncover further introgression patterns among species within *Camellia,* an economically important genus of Theaceae.

### Whole genome duplication

Two rounds of ancient WGD events, i.e. Ad-γ and Ad-β, occurred in the tea tree *(Camellia sinensis* var. *assamica)* genome (Xia *et al.*, 2017; Xia *et al.*, 2020). However, analysis of genic collinearity reveals that a recent WGD event occurred after the divergence of tea and kiwifruit lineages, based on the genome of another variety of tea *(Camellia sinensis* var. *sinensis)* (Wei *et al.*, 2018), Larson *et al.* (2020) named this WGD as Cm-α. Recently, based on a chromosome-scale genome of *Camellia sinensis* var. *sinensis,* the authors suggested one recent *Camellia* tetraploidization event occurred after the divergence of *C. sinensis* and *A. chinensis* from their common ancestor (Chen *et al.*, 2020). But the time of the *Camellia* tetraploidization event (58.9-61.7 Ma) was very close to the divergence time between *C. sinensis* and *A. chinensis* at 61.2-65.3 Ma. Here, we identified a WGD event shared by all genera within Theaceae, and also other families such as Symplocaceae, Styracaceae, Pentaphylacaceae and Diapensiaceae (Fig. 4, Fig. S4), and also clarified that Cm-α proposed by Wei *et al.* (2018) and Larson *et al.* (2020) and the recent tetraploidization event found by Chen *et al.* (2020) were actually Ad-β (i.e. Ericales clade). Ad-β has been recently revised to the core Ericales clade according to genome collinearity and also MAPS pipeline (Leebens-Mack *et al.*, 2019), and more specifically to the core Ericales+Primuloids+Polemonioids+Lecythidaceae clade, using deep Asterid phylotranscriptomic analyses (Zhang *et al.*, 2020). Thus, our results support that the tea family experienced two WGD events in the evolutionary history, i.e. Ad-γ and Ad-β, no other specific WGD event occurred within the family.

## Acknowledgement

This work was supported by National Natural Science Foundation of China (No. 32070369, 31700182), the Large-scale Scientific Facilities of the Chinese Academy of Sciences (No. 2017-LSFGBOWS-02), open Research Fund of Guangxi Key Laboratory of Special Non-wood Forest Cultivation & Utilization (No. 19-B-01-03), the Youth Innovation Promotion Association CAS (No. 2021393) and CAS “Light of West China” Program. The authors are grateful to Prof. Liang Fang (Jiujiang Unversity), Prof. Zhong-Lang Wang, Drs. Jie Cai, Ting Zhang, Yun-Long Liu (Kunming Institute of Botany, Chinese Academy of Sciences) for their help with sample collection and data analysis, and to Mark Whitten, Sheng-Chen Shan (University of Florida) for their assistance for sampling of *Stewartia malacodendron.*

## Notes

### Competing Interest Statement

The authors have declared no competing interest.

